# Genetic data and cognitively-defined late-onset Alzheimer’s disease subgroups

**DOI:** 10.1101/367615

**Authors:** Shubhabrata Mukherjee, Jesse Mez, Emily Trittschuh, Andrew J. Saykin, Laura E. Gibbons, David W. Fardo, Madeline Wessels, Julianna Bauman, Mackenzie Moore, Seo-Eun Choi, Alden L. Gross, Joanne Rich, Diana K.N. Louden, R. Elizabeth Sanders, Thomas J. Grabowski, Thomas D. Bird, Susan M. McCurry, Beth E. Snitz, M. Ilyas Kamboh, Oscar L. Lopez, Philip L. De Jager, David A. Bennett, C. Dirk Keene, Eric B. Larson, Paul K. Crane

## Abstract

Categorizing people with late-onset Alzheimer’s disease into biologically coherent subgroups is important for personalized medicine. We evaluated data from five studies (total n=4 050, of whom 2 431 had genome-wide single nucleotide polymorphism (SNP) data). We assigned people to cognitively-defined subgroups on the basis of relative performance in memory, executive functioning, visuospatial functioning, and language at the time of Alzheimer’s disease diagnosis. We compared genotype frequencies for each subgroup to those from cognitively normal elderly controls. We focused on *APOE* and on SNPs with p<10^-5^ and odds ratios more extreme than those previously reported for Alzheimer’s disease (<0.77 or >1.30). There was substantial variation across studies in the proportions of people in each subgroup. In each study, higher proportions of people with isolated substantial relative memory impairment had ≥1 *APOE* ε4 allele than any other subgroup (overall p= 1.5 × 10^-27^). Across subgroups, there were 33 novel suggestive loci across the genome with p<10^-5^ and an extreme OR compared to controls, of which none had statistical evidence of heterogeneity and 30 had ORs in the same direction across all datasets. These data support the biological coherence of cognitively-defined subgroups and nominate novel genetic loci.

## Introduction

Clinical heterogeneity is common among people with late-onset Alzheimer’s disease; see^1^ for a review. Categorizing people with a condition into biologically coherent subgroups is an important personalized medicine strategy.^2^ This strategy is particularly recommended for neurodegenerative conditions.^3^ Once biologically coherent subgroups are identified, further investigations may elucidate subgroup-specific treatments.

Genetic data may be useful for determining whether a proposed categorization strategy results in biologically coherent subgroups; see the Box.

We have developed an approach for categorizing people with late-onset Alzheimer’s disease based on relative performance across cognitive domains. We determine each person’s average performance at diagnosis across memory, executive functioning, language, and visuospatial ability, and consider relative impairments in each domain from that average. We previously evaluated one study’s data and showed that our strategy identified a subgroup with higher degrees of amyloid angiopathy and higher proportions with ≥1 *APOE* ε4 allele.^4^

Here we evaluate data from five studies with people with late-onset Alzheimer’s disease^5^ and cognitively normal elderly controls.^6^ We used modern psychometric approaches to co-calibrate cognitive scores. We used scores to identify subgroups.

We used genetic data to determine whether our categorization identifies biologically coherent subgroups.

## Materials and Methods

### Study design and participants

We used data from the Adult Changes in Thought (ACT) study, the Alzheimer’s Disease Neuroimaging Initiative (ADNI), the Rush Memory and Aging Project (MAP) and Religious Orders Study (ROS), and the University of Pittsburgh Alzheimer Disease Research Center (PITT). Each study has published widely, and their genetic data are included in large analyses of late-onset Alzheimer’s disease.^6, 7^ All five studies use the same research criteria to define clinical Alzheimer’s disease.^5^ Three studies (ACT, MAP, and ROS) are prospective cohort studies that enroll cognitively normal elderly individuals and follow them to identify incident dementia cases. For these, we analyzed cognitive data from the visit with the incident clinical Alzheimer’s disease diagnosis. Two studies (ADNI and PITT) are clinic-based research cohort studies. For those studies we analyzed cognitive data from the first study visit for people with prevalent Alzheimer’s disease; we limited inclusion to those with Clinical Dementia Rating Scale^8^ of 0.5 or 1, indicating mild Alzheimer’s disease. For people from ADNI or PITT who did not initially have Alzheimer’s disease, we analyzed cognitive data from the incident Alzheimer’s disease visit.

In each study, we included people diagnosed with late-onset Alzheimer’s disease as defined by research criteria.^5^ We used data from everyone for all analyses other than genetic analyses; we limited those to self-reported whites. For genetic analyses we also used data from self-reported white cognitively normal elderly controls from each study. Details on those cohorts are included in reports from the parent studies and in Supplemental Text 6 (derived from Lambert et al.^6^).

### Cognitive data procedures

Staff from each study administered a comprehensive neuropsychological test battery that included assessment of memory, executive functioning, language, and visuospatial functioning. We obtained granular (“item-level”) data from each parent study. Each stimulus administered to a participant was deemed an “item”. As outlined in our previous paper^2^, every item administered in each study was considered by our panel of experts (JM, ET, AJS). Our panel designated each item as primarily a measure of memory, executive functioning, language, visuospatial functioning, or none of these. They also assigned items to theory-driven subdomains.

We carefully considered items where the same stimulus was administered to participants across different studies to identify “anchor items” that could anchor metrics across studies. We reviewed response categories recorded by each study for these overlapping items to ensure that consistent scoring was used across studies. We identified anchor items as those overlapping items with identical scoring across studies. We used bifactor models in Mplus^9^ to co-calibrate separate scores for memory, executive functioning, language, and visuospatial functioning.

Details of item assignments, psychometric analyses in each study, and co-calibration methodology across studies are provided in Supplemental Texts 1, 2, and 5. Code is available from the authors upon request.

We used the ACT sample of people with incident Alzheimer’s disease as our reference population for the purpose of scaling domain scores; ACT was our largest sample from a prospective cohort study of people with late-onset Alzheimer’s disease. We z-transformed scores from other studies to ACT-defined metrics for each domain.

## Assignment to subgroups

We assigned people to subgroups as we have done previously.^4^ Briefly, for each person we determined their average score across memory, executive functioning, language, and visuospatial functioning. We determined differences between each domain score and this average score. We identified domains with substantial relative impairments as those with relative impairments at least as large as 0.80 standard deviation units as explained in Supplemental Text 3. We categorized people by the number of domains with substantial relative impairments (0, 1, or ≥2) and further categorized those with substantial relative impairments in a single domain by the domain with a substantial relative impairment. This approach results in six potential subgroups: those with no domain with a substantial relative impairment; those with an isolated substantial relative impairment in one of four domains (e.g., isolated substantial relative memory impairment, isolated substantial relative language impairment, etc.), and those with multiple domains with substantial relative impairments.

## Statistical analyses

As in our previous study,^4^ we compared the proportion of people with late-onset Alzheimer’s disease in each subgroup with at least one *APOE* ε4 allele. For other genetic analyses we combined data from ROS and MAP, as has been done many times previously, and evaluated data separately in four genetic datasets. Each dataset was imputed using IMPUTE2 and samples of European ancestry from the 2012 build of the 1000 Genomes project. We excluded SNPs with R^2^ or information scores < 0.5, and SNPs with a minor allele frequency <3%. Further details are provided in Supplemental Text 6 and in Lambert et al.^6^

We used KING-Robust^10^ to obtain study-specific principal components to account for population stratification. We used logistic regression in PLINK v 1.9^11^ to evaluate associations at each SNP for each cognitively-defined subgroup. Cognitively normal elderly controls from each study were the comparison group for all of these analyses. We included covariates for age, sex, and principal components. We used METAL^12^ for meta-analysis.

IGAP’s most extreme odds ratio (OR) outside of chromosome 19 was for rs11218343 associated with *SORL1*, which had an OR of 0.77 reported in the Stage 1 and 2 meta-analysis.^6^ We focused attention on SNPs where meta-analysis ORs for any cognitively-defined subgroup were <0.77 (or, equivalently, ORs >[1/0.77], which is >1.30). As presented in the Box, more extreme ORs in a single subgroup, with strong replication across genetic datasets, would represent strong support of biologically coherent categorization.

For genetic loci previously identified as associated with risk of late-onset Alzheimer’s disease, we used the methods described in^13^ applied to IGAP’s previously reported ORs and confidence intervals to determine significance of subgroup associations compared to IGAP.

We used genetic data from all cognitively normal elderly controls and all people with Alzheimer’s disease to generate Alzheimer’s disease and subgroup genetic risk scores. We used 1) IGAP SNPs and effect sizes to generate Alzheimer’s disease risk scores, and 2) our results to generate risk scores for each of the five subgroups.

To evaluate these risk scores, we used logistic regression with Alzheimer’s disease case vs. control status as the outcome, and included terms for age and sex. We compared a model with just the addition of the IGAP genetic risk score to a model that incorporated that score plus the five subgroup risk scores. Finally, we compared area under the receiver operator characteristic (ROC) curves for the model with the IGAP risk score to a model that did not include that term but included terms for the five subgroup risk scores. Further details are shown in Supplemental Text 12.

## Data sharing

Co-calibrated scores for each domain are available from the parent studies. GWAS meta-analysis summary statistics for each subgroup will be available on the National Institute on Aging Genetics of Alzheimer’s Disease Storage Site (NIAGADS; www.niagads.org).

## Results

In all, there were 4 050 people with sufficient cognitive data to be classified into a subgroup. Demographic characteristics and average cognitive domain scores by study are shown in Table 1. Participants in the prospective cohort studies (ACT, MAP, and ROS) were older on average than those from ADNI and PITT. Most participants in each study self-reported white race (90% in ACT to 96% in MAP). There was some variation in cognitive performance across studies. The most notable differences from ACT (our reference for scaling) were for executive functioning in ADNI and ROS (average 0.8 units higher), and for language in MAP (average 0.8 units lower).

**Table 1.** Demographic and cognitive characteristics of people with late-onset Alzheimer’s disease by study and overall.

| **Characteristic** | **ACT**<br>**(n=825)** | **ADNI**<br>**(n=650)** | **MAP**<br>**(n=404)** | **ROS**<br>**(n=419)** | **PITT**<br>**(n=1 752)** | **Overall**<br>**(n=4 050)** |
| --- | --- | --- | --- | --- | --- | --- |
| Female sex, n (%) | 522 (63%) | 267 (41%) | 285 (71%) | 302 (72%) | 1 106 (63%) | 2 482 (61%) |
| Age at diagnosis, mean (SD) | 86 (6) | 77 (7) | 87 (6) | 85 (6) | 76 (7) | 80 (8) |
| Years of education, mean (SD) | 14 (3) | 15 (3) | 14 (3) | 18 (3) | 14 (3) | 14 (3) |
| Self-reported white race, n (%) | 743 (90%) | 613 (94%) | 386 (96%) | 382 (91%) | 1 595 (91%) | 3 719 (92%) |
| Composite scores for each cognitive domain, mean (SD) |  |  |  |  |  |  |
| Memory | 0.0 (1.0) | 0.0 (1.0) | -0.5 (1.1) | 0.0 (1.0) | -0.6 (1.1) | -0.3 (1.1) |
| Visuospatial | 0.0 (1.0) | 0.3 (1.0) | 0.1 (1.1) | 0.1 (1.0) | 0.4 (1.5) | 0.3 (1.3) |
| Executive function | 0.0 (1.0) | 0.8 (1.2) | 0.5 (1.0) | 0.8 (1.0) | 0.3 (0.8) | 0.4 (1.0) |
| Language | 0.0 (1.0) | 0.5 (1.0) | -0.8 (1.2) | -0.4 (1.1) | -0.1 (1.2) | -0.1 (1.2) |

Proportions of people in each subgroup are shown in Figure 1. There was considerable heterogeneity in proportions across studies (*χ*^2^_df=20_ = 468.7, p=1.0×10–86).

**Figure 1.**
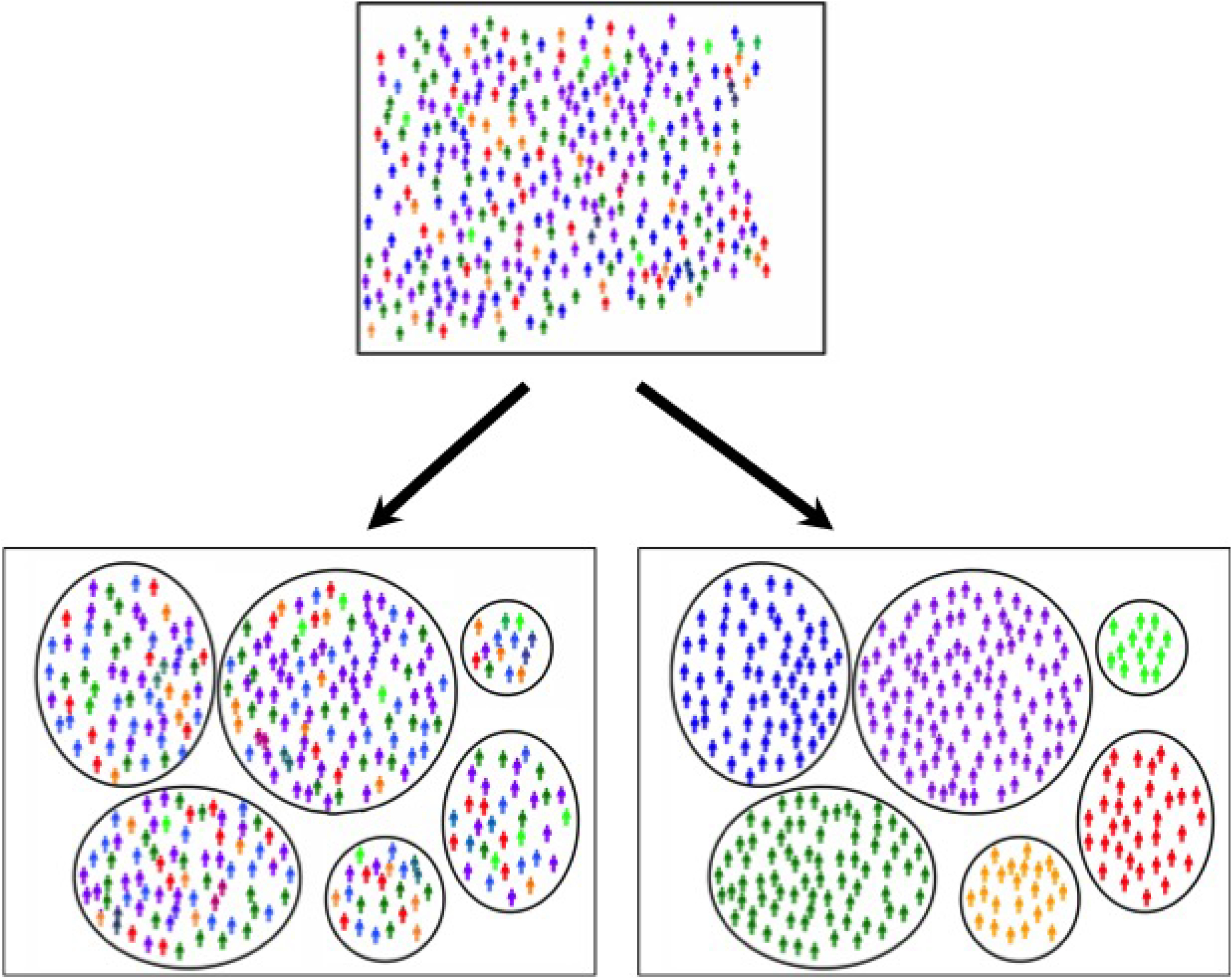
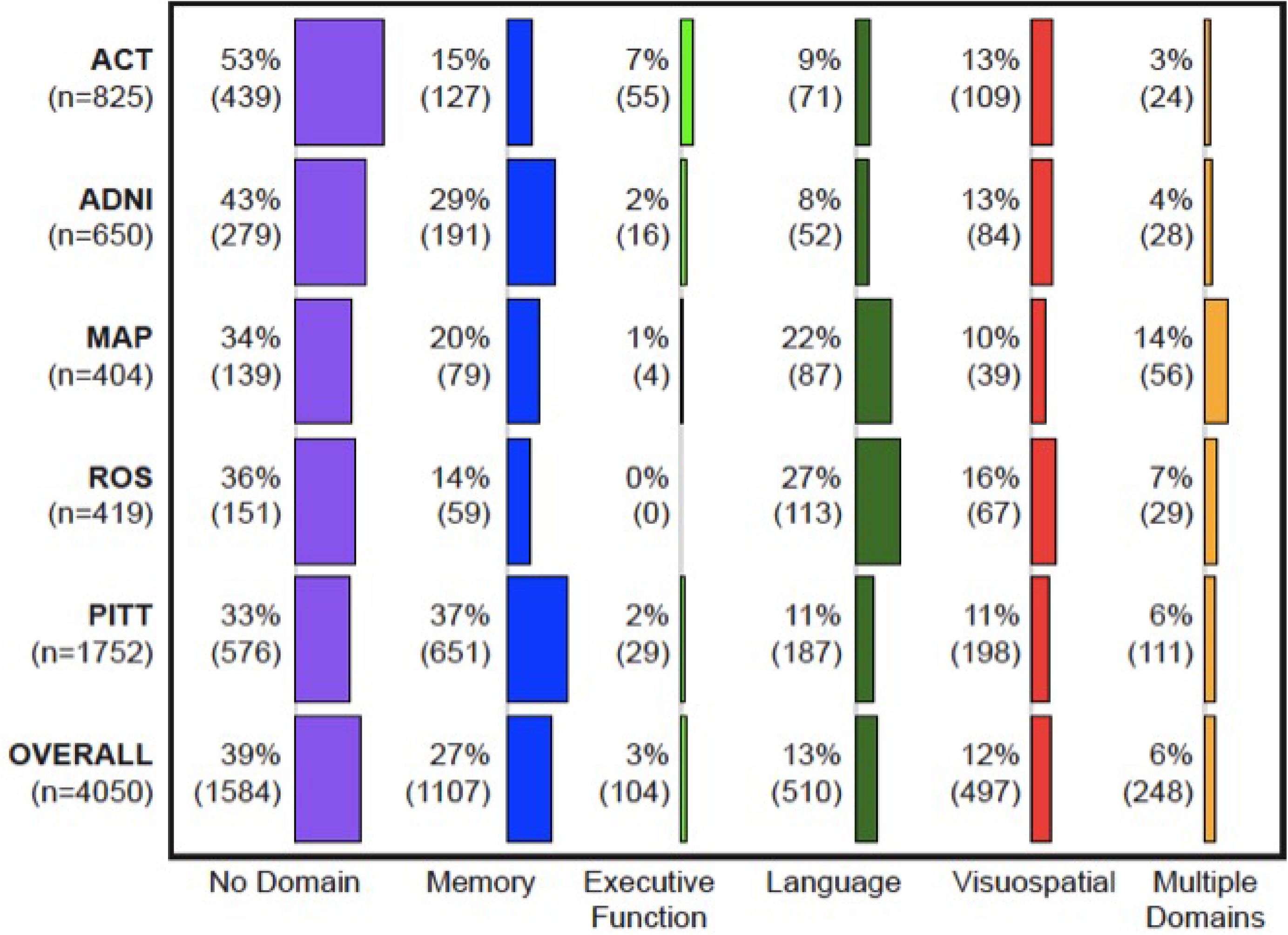
Proportions of people in each study and overall in each cognitively-defined subgroup.

Demographic characteristics of people in each subgroup were similar to those for people with Alzheimer’s disease overall (Supplemental Table 29). The proportion who were female ranged from 51% for isolated substantial relative executive functioning impairment to 63% of those with isolated substantial relative visuospatial impairment. Mean age at diagnosis ranged from 79 for those with isolated substantial relative visuospatial or memory impairment to 82 for those with isolated substantial relative language impairment. Mean years of education did not vary substantially across subgroups.

There were 3 701 people with *APOE* genotype data (Table 2). We published *APOE* results from ACT^4^; the proportion of those with isolated substantial memory impairment with ≥1 *APOE* ε4 allele was 12% higher than overall in that study. This finding was consistent across all five studies. Overall, the proportion of people with ≥ 1 *APOE*e4 allele was 15% higher those with isolated substantial memory impairment (65%) compared with the entire sample (50%). The differences in proportions with ≥1 *APOE* ε4 allele were highly significant (p=1.5×10^-27^). The *APOE* result was not sensitive to choosing other thresholds to indicate a substantial relative impairment (Supplemental Text 4 and Supplemental Figure 2).

There were 2 431 people with late-onset Alzheimer’s disease and 3447 cognitively normal elderly controls with genome-wide SNP data. Top results are shown in Figure 2. There were 33 loci outside Chromosome 19 where the p value for one subgroup was <5×10^-5^. All of these had ORs <0.77 or >1.30 compared to cognitively normal elderly controls. These included nine loci for those with isolated substantial visuospatial impairment (red dots, including rs2289506 near *NIT2* on chromosome 3, rs9369477 near *SPATS1* on chromosome 4, rs2046197 near *CSMD1* on chromosome 8, and rs8091629 near *SLC14A2* on chromosome 18), nine for those with multiple domains with substantial relative impairments (yellow dots, including rs698842 near *NRXN1*on chromosome 2, rs78872508 near *HDAC9* on chromosome 7, and rs4348488 near *BMP1*on chromosome 8), seven for those with no domain with a substantial relative impairment (purple dots, including rs11708767 near *MED12L* on chromosome 3, rs72839770 near *DVL2* on chromosome 17, and rs7264688 near *MGME1* on chromosome 20), six for those with isolated substantial language impairment (green dots, including rs13374908 near *FAM163A* on chromosome 1, rs28715896 near *ERBB4*on chromosome 2, and rs75337321 near *CACNA2D3* on chromosome 3), and two for those with isolated substantial memory impairment (blue dots, including rs1977412 near *AGT*on chromosome 1), and one for those with no domains with substantial impairments (rs7264688 near *MGME1* on chromosome 20). Other loci shown in Figure 2 not named in the preceding sentence were >50kb from genes. Replication results are in Supplemental Table 2a and 2b. All ORs were in the same direction for all these SNPs except rs28715896 on chromosome 2 near *ERBB4*, rs61835453 on chromosome 10, and rs365521 on chromosome 17. Heterogeneity p values did not suggest heterogeneity for any of these SNPs. No SNP outside the *APOE* region reached p<5×10^-8^, the traditional level of genome-wide significance (Figure 2).

**Figure 2.**
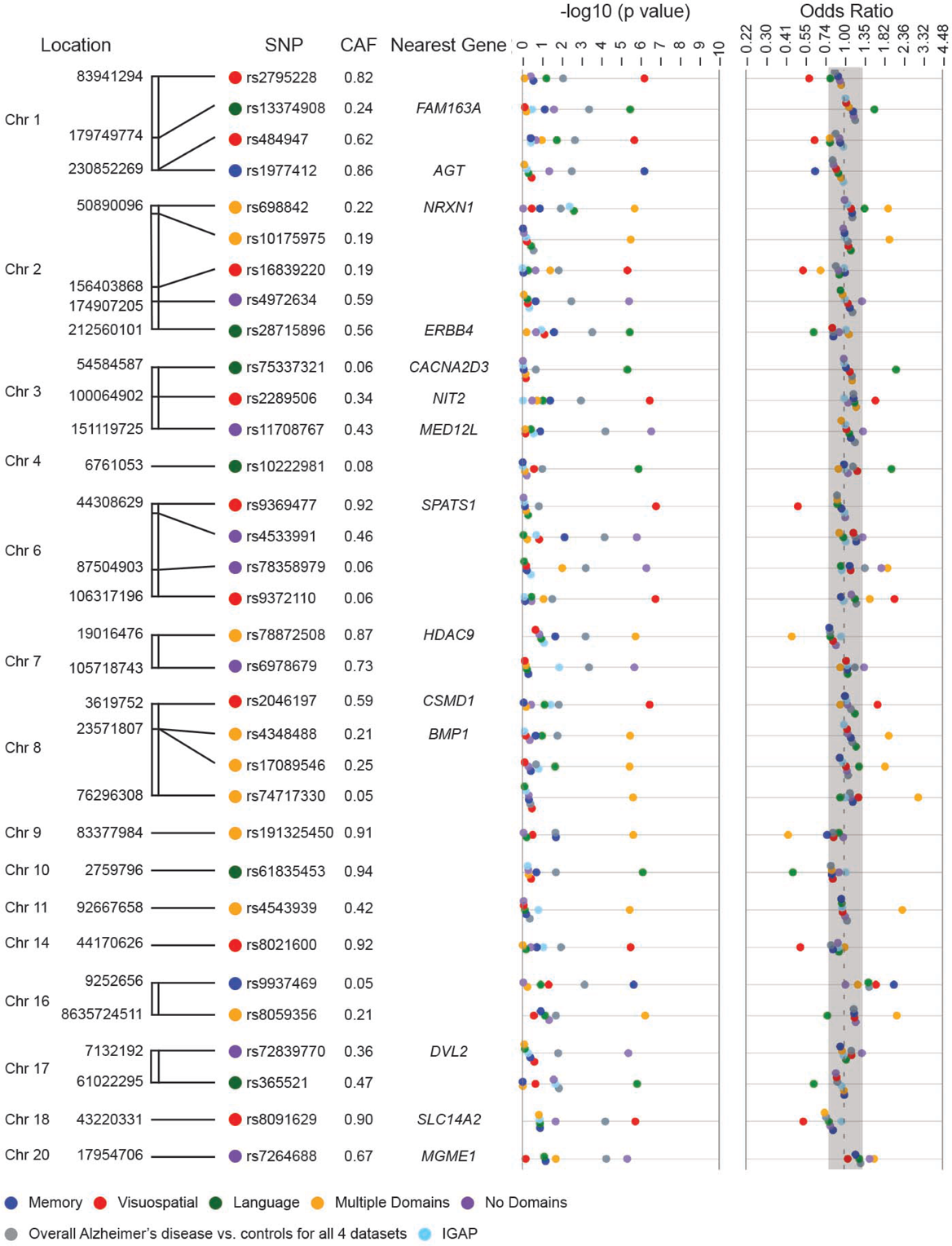
Novel SNPs associated with cognitively defined subdomains with p<10–5 and OR<0.77 or >1.33^1^. 1. CAF = coded allele frequency. SNPs further than 50 kB from a gene do not have a gene name reported here. Gray shading in the odds ratios column of the figure delineates ORs>0.77 and <1.30, which is the range of ORs outside *APOE* from the International Genomics of Alzheimer’s Project (IGAP)^6^

**Table 2.** Proportion of those in each cognitively defined subgroup with ≥1 *APOE* ϵ4 allele, by study and overall.

| **Study** | **No Domain** | **Memory** | **Executive** | **Language** | **Visuospatial** | **Multiple Domains** | **Overall** | **p-value<sup>1</sup>** |
| --- | --- | --- | --- | --- | --- | --- | --- | --- |
| ACT (n=711) | 134 (35%) | 51 (46%) | 11 (24%) | 14 (24%) | 27 (30%) | 6 (32%) | 243 (34%) | 0.022 |
| ADNI (n=650) | 185 (66%) | 139 (73%) | 10 (63%) | 30 (58%) | 48 (57%) | 15 (54%) | 427 (66%) | 0.07 |
| MAP (n=386) | 45 (34%) | 33 (44%) | 1 (33%) | 23 (27%) | 9 (26%) | 21 (39%) | 132 (34%) | 0.24 |
| ROS (n=393) | 49 (35%) | 26 (48%) | — | 31 (28%) | 26 (41%) | 11 (46%) | 143 (36%) | 0.08 |
| PITT (n=1 561) | 283 (54%) | 411 (70%) | 15 (65%) | 72 (44%) | 97 (57%) | 39 (44%) | 917 (59%) | $5.1 \times 10^{-11}$ |
| Overall (n=3 701) | 696 (48%) | 660 (65%) | 37 (42%) | 170 (36%) | 207 (47%) | 92 (43%) | 1 861 (50%) | $1.5 \times 10^{-27}$ |

1. p values based on $\chi^2_{(4)}$ for ROS, otherwise $\chi^2_{(5)}$

1. p values based on χ^2^_(4)_ for ROS, otherwise χ^2^_(5)_

Results for selected IGAP loci are shown in Supplemental Table 30. We selected subgroups with meta-analytic ORs <0.77 or >1.30 and for which results from all four data sets were in the same direction. Results for other IGAP SNPs for all studies and all subgroups are in Supplemental Table 31.

Across all datasets, there were 5 878 people with SNP data who were either normal controls or Alzheimer’s disease cases. The logistic regression model for case vs. control status with age, sex, and IGAP gene scores had a likelihood of 176.13 and a pseudo-R^2^ of 0.022. A model that included all of those terms and also included five subgroup gene scores had a likelihood of 311.94 and a pseudo-R^2^ of 0.039. These were nested models; we compared them with a likelihood ratio test with 5 degrees of freedom. The difference in likelihoods was highly significant (p=1.39×10^-27^).

The area under the ROC curve for the model with age, sex, and the IGAP genetic risk score was 0.60 (95% CI 0.58, 0.61), while for the model with age, sex, and five subgroup genetic risk scores, the area under the ROC curve was 0.62 (95% CI 0.61, 0.64). This difference was statistically significant (*χ*^2^_df=1_ = 11.15, p=0.0008). Further analyses are described in Supplemental Text 12.

## Discussion

We used modern psychometric methods to co-calibrate cognitive data to generate domain scores across five different studies of older adults with research-quality clinical Alzheimer’s disease diagnoses. We obtained scores on the same metric, so scores from different studies were directly comparable to each other. The proportion of people with Alzheimer’s disease in each study categorized in each subgroup varied across studies (Figure 1). We used genetic data to determine whether our categorization scheme resulted in biologically coherent subgroups. Top genetic associations from each subgroup were consistent across studies, suggesting our findings were not due to idiosyncrasies of any particular study. Gene scores for subgroups performed better in predicting case vs. control status than gene scores derived from Alzheimer’s disease case-control analyses.

*APOE* genotype was significantly different across subgroups. We previously showed in ACT that more people with isolated substantial relative memory impairment had at least one *APOE* ε4 allele than people in other subgroups^4^. We robustly replicated that finding here (Table 2). Associations between *APOE*e4 alleles and memory impairment among people with Alzheimer’s disease have been previously noted^14^.

Outside of chromosome 19 which was dominated by *APOE*-related signals for all subgroups, we identified 33 loci with p<10^-5^ for at least one subgroup. All of these had ORs <0.77 or >1.30 (Figure 2). There were consistent findings across datasets for nearly all these loci (Supplemental Text 7–11).

Gene scores from IGAP SNPs explained a modest amount of risk for case-control status. Adding gene scores from our cognitively defined subgroups improved prediction of Alzheimer’s disease. Furthermore, in a head-to-head comparison, gene scores for cognitively defined subgroups did a better job predicting Alzheimer’s disease status than did gene scores from IGAP SNPs.

These data provide strong support for the biological coherence of subgroups produced by our categorization scheme. Each subgroup we analyzed has extreme ORs at novel SNPs that were consistent across multiple independent samples. Even with the relatively small sample sizes from these studies, the large effect sizes at common SNPs produced p values that are close to genome-wide significance.

Others have used different data sources to categorize people with Alzheimer’s disease. Sweet and colleagues compared people with Alzheimer’s disease who developed psychosis to those who did not.^15^ They identified a few interesting loci, but effect sizes were much smaller than those reported here (Supplemental Text 13).

We were part of a consortium evaluating rates of decline among people with Alzheimer’s disease.^16^ The evidence in support of rates of decline, as an organizing characteristic among people with Alzheimer’s disease, is not nearly as strong as that shown here for cognitively-defined subgroups.

Others have used cluster analysis approaches applied to neuropsychological^17, 18^ or imaging^19, 20^ data to categorize people with Alzheimer’s disease. There are very important distinctions between those approaches and the approach adopted here. In cluster analysis, the computer maximizes some distance across groups in a way that may not make clinical or biological sense. Disease severity is an important consideration; see^21^ for a nice discussion.

Our approach began with theory and focused exclusively on cognitive data. An early paper considered differences between memory and executive functioning among people with Alzheimer’s disease.^22^ Differences between these scores enables memory to serve as something of a proxy for disease severity. This framework is useful for considering dysexecutive Alzheimer’s disease.^23–26^

We have extended that framework to incorporate additional cognitive domains. The field has increasingly emphasized the importance of Alzheimer’s disease variants including primary progressive aphasia (PPA) and posterior cortical atrophy (PCA);^27^ these rare subtypes are described as typically having early onset. Clinical descriptions of the cognitive patterns of these variants emphasize relative deficits between language (PPA) or visuospatial functioning (PCA) and other domains. We thus incorporate average performance across domains, and differences from that average, to more fully capture the range of clinical heterogeneity described in late onset Alzheimer’s disease.^1^

Our results should be considered mindful of limitations of our study. Data evaluated here are from studies with well-educated people of European ancestry. It will be important to replicate this approach among people with diverse genetic backgrounds. While we combined data from five large studies, the resulting subgroups were underpowered to reach genome-wide significance, and one subgroup (isolated substantial relative executive functioning impairment) was too small to analyze at all. It will be important to incorporate additional data sets to see whether novel suggestive loci reach genome-wide significance, and to identify additional loci. We used a large threshold of 0.80 SD to characterize “substantial” relative impairments, which may be too conservative. Our categorization approach relies exclusively on cognitive data. We could imagine a more optimal approach that also incorporates imaging and/or fluid biomarkers.

In conclusion, genome-wide genetic data enabled us to determine that a cognitively-defined categorization scheme produced biologically coherent subgroups of people with Alzheimer’s disease. This is an important result on the road towards personalized medicine.

## Acknowledgements

Analyses were funded by R01 AG042437 (P Crane, PI) and R01 AG029672 (P Crane, PI).

Data archiving was supported by R01 AG042437-S1 (P Crane, PI)

Dr. Mez’s efforts were also supported by K23 AG046377 (J Mez, PI)

The efforts of Dr. Gibbons, Dr. Grabowski, and Dr. Keene were also supported by P50 AG005136 (T Grabowski, PI). Dr. Keene’s efforts were also supported by the Nancy and Buster Alvord Endowment.

Dr. Gross’s efforts were supported by K01 AG050699 (A Gross, PI).

Dr. Bird’s efforts were supported by Department of Veterans Affairs Research Funds.

Dr. Saykin’s efforts were supported by U01 AG042904 (M Weiner, PI), P30 AG010133 (A Saykin, PI), and R01 AG019771 (A Saykin, PI).

ACT data collection was supported by U01 AG006781 (E Larson and P Crane, MPIs). ACT genotyping was supported by U01 HG006375 (E Larson and G Jarvik, MPIs). Cerebellum samples for genotyping for some ACT samples were prepared with support from P50 AG005136 (T Grabowski, PI). The authors thank Aimee Schantz and Allison Beller for administrative support for the ACT cerebellum samples.

ADNI data collection and genotyping were supported by U01 AG024904 (M Weiner, PI). Data collection and sharing for this project was funded by the Alzheimer’s Disease Neuroimaging Initiative (ADNI) (National Institutes of Health Grant U01 AG024904) and DOD ADNI (Department of Defense award number W81XWH-12–2-0012). ADNI is funded by the National Institute on Aging, the National Institute of Biomedical Imaging and Bioengineering, and through generous contributions from the following: AbbVie, Alzheimer’s Association; Alzheimer’s Drug Discovery Foundation; Araclon Biotech; BioClinica, Inc.; Biogen; Bristol-Myers Squibb Company; CereSpir, Inc.; Cogstate; Eisai Inc.; Elan Pharmaceuticals, Inc.; Eli Lilly and Company; EuroImmun; F. Hoffmann-La Roche Ltd and its affiliated company Genentech, Inc.; Fujirebio; GE Healthcare; IXICO Ltd.; Janssen Alzheimer Immunotherapy Research & Development, LLC.; Johnson & Johnson Pharmaceutical Research & Development LLC.; Lumosity; Lundbeck; Merck & Co., Inc.; Meso Scale Diagnostics, LLC.; NeuroRx Research; Neurotrack Technologies; Novartis Pharmaceuticals Corporation; Pfizer Inc.; Piramal Imaging; Servier; Takeda Pharmaceutical Company; and Transition Therapeutics. The Canadian Institutes of Health Research is providing funds to support ADNI clinical sites in Canada. Private sector contributions are facilitated by the Foundation for the National Institutes of Health (www.fnih.org). The grantee organization is the Northern California Institute for Research and Education, and the study is coordinated by the Alzheimer’s Therapeutic Research Institute at the University of Southern California. ADNI data are disseminated by the Laboratory for Neuro Imaging at the University of Southern California.

MAP data collection and genotyping were supported by R01 AG017917 (D Bennett, PI). Additional genotyping was supported by Kronos, Zinfandel, and U01 AG032984 (G Schellenberg, PI).

ROS data collection and genotyping were supported by P30 AG10161 (D Bennett, PI), R01 AG15819 (D Bennett, PI), and R01 AG30146 (D Evans, PI). Additional genotyping was supported by Kronos, Zinfandel, and U01 AG032984 (G Schellenberg, PI).

PITT data collection were funded by P50 AG05133 (O Lopez PI), R01 AG030653 (MI Kamboh, PI), and R01 AG041718 (MI Kamboh, PI).

## Supplemental information

Supplementary information is available at MP’s website

## Figure Legends

### Box. Schematic representation of incoherent vs. coherent subgrouping

The large group at the top represents a heterogeneous group of individuals. A strategy is applied to categorize individuals into subgroups. For a precision medicine approach to work, the categorization should reduce heterogeneity. In the lower left figure, the method did not reduce heterogeneity and thus, we refer to this as an *incoherent subgrouping strategy*. In contrast, the lower right figure was produced by a different method which resulted in relatively homogenous subgroups; this method would represent a *coherent subgrouping strategy*.

For incoherent subgroup comparisons with controls, top genetic hits and effect sizes would not be expected to be different than those observed in the entire group. Further, for a given incoherent subgroup, spurious genetic associations at a locus would not be expected to replicate in that subgroup in other datasets. In contrast, for coherent subgroup comparisons with controls, there is improved potential for identification of novel loci and effect sizes could be stronger than those seen for the original ungrouped data. Further replication of subgroup associations in other datasets would occur more often than expected by chance.

Genetic data may serve a useful role in determining whether a categorization strategy produces biologically coherent subgroups.

## References

1. Lam B, Masellis M, Freedman M, Stuss DT, Black SE. Clinical, imaging, and pathological heterogeneity of the Alzheimer’s disease syndrome. Alzheimers Res Ther 2013; 5(1): 1.

2. Cholerton B, Larson EB, Quinn JF, Zabetian CP, Mata IF, Keene CD et al. Precision medicine: clarity for the complexity of dementia. Am J Pathol 2016; 186(3): 500–506.

3. Girard SL, Rouleau GA. Genome-wide association study in FTD: divide to conquer. Lancet Neurol 2014; 13(7): 643–644.

4. Crane PK, Trittschuh E, Mukherjee S, Saykin AJ, Sanders RE, Larson EB et al. Incidence of cognitively defined late-onset Alzheimer’s dementia subgroups from a prospective cohort study. Alzheimer’s & Dementia 2017; 13(12): 1307–1316.

5. McKhann G, Drachman D, Folstein M, Katzman R, Price D, Stadlan EM. Clinical diagnosis of Alzheimer’s disease: report of the NINCDS-ADRDA Work Group under the auspices of Department of Health and Human Services Task Force on Alzheimer’s Disease. Neurology 1984; 34(7): 939–944.

6. Lambert JC, Ibrahim-Verbaas CA, Harold D, Naj AC, Sims R, Bellenguez C et al. Meta-analysis of 74,046 individuals identifies 11 new susceptibility loci for Alzheimer’s disease. Nat Genet 2013 45(12): 1452–1458.

7. Naj AC, Jun G, Beecham GW, Wang LS, Vardarajan BN, Buros J et al. Common variants at MS4A4/MS4A6E, CD2AP, CD33 and EPHA1 are associated with late-onset Alzheimer’s disease. Nat Genet 2011 43(5): 436–441.

8. Morris JC. The Clinical Dementia Rating (CDR): current version and scoring rules. Neurology 1993; 43(11): 2412–2414.

9. Muthen LK, Muthen BO. Mplus user’s guide. 7 edn. Muthen & Muthen: LA, 1998–2012.

10. Manichaikul A, Mychaleckyj JC, Rich SS, Daly K, Sale M, Chen WM. Robust relationship inference in genome-wide association studies. Bioinformatics 2010; 26(22): 2867–2873.

11. Chang CC, Chow CC, Tellier LC, Vattikuti S, Purcell SM, Lee JJ. Second-generation PLINK: rising to the challenge of larger and richer datasets. Gigascience 2015; 4: 7.

12. Willer CJ, Li Y, Abecasis GR. METAL: fast and efficient meta-analysis of genomewide association scans. Bioinformatics 2010; 26(17): 2190–2191.

13. Schott JM, Crutch SJ, Carrasquillo MM, Uphill J, Shakespeare TJ, Ryan NS et al. Genetic risk factors for the posterior cortical atrophy variant of Alzheimer’s disease. Alzheimers Dement 2016; 12(8): 862–871.

14. El Haj M, Antoine P, Amouyel P, Lambert JC, Pasquier F, Kapogiannis D. Apolipoprotein E (APOE) epsilon4 and episodic memory decline in Alzheimer’s disease: A review. Ageing Res Rev 2016; 27: 15–22.

15. DeMichele-Sweet MAA, Weamer EA, Klei L, Vrana DT, Hollingshead DJ, Seltman HJ et al. Genetic risk for schizophrenia and psychosis in Alzheimer disease. Mol Psychiatry 2017.

16. Sherva R, Tripodis Y, Bennett DA, Chibnik LB, Crane PK, de Jager PL et al. Genome-wide association study of the rate of cognitive decline in Alzheimer’s disease. Alzheimers Dement 2014; 10(1): 45–52.

17. Scheltens NM, Galindo-Garre F, Pijnenburg YA, van der Vlies AE, Smits LL, Koene T et al. The identification of cognitive subtypes in Alzheimer’s disease dementia using latent class analysis. J Neurol Neurosurg Psychiatry 2016; 87(3): 235–243.

18. Scheltens NME, Tijms BM, Koene T, Barkhof F, Teunissen CE, Wolfsgruber S et al. Cognitive subtypes of probable Alzheimer’s disease robustly identified in four cohorts. Alzheimers Dement 2017.

19. Noh Y, Jeon S, Lee JM, Seo SW, Kim GH, Cho H et al. Anatomical heterogeneity of Alzheimer disease: based on cortical thickness on MRIs. Neurology 2014; 83(21): 1936–1944.

20. Dong A, Toledo JB, Honnorat N, Doshi J, Varol E, Sotiras A et al. Heterogeneity of neuroanatomical patterns in prodromal Alzheimer’s disease: links to cognition, progression and biomarkers. Brain 2017; 140(3): 735–747.

21. Ossenkoppele R, Cohn-Sheehy BI, La Joie R, Vogel JW, Moller C, Lehmann M et al. Atrophy patterns in early clinical stages across distinct phenotypes of Alzheimer’s disease. Hum Brain Mapp 2015; 36(11): 4421–4437.

22. Dickerson BC, Wolk DA. Dysexecutive versus amnesic phenotypes of very mild Alzheimer’s disease are associated with distinct clinical, genetic and cortical thinning characteristics. J Neurol Neurosurg Psychiatry 2011; 82(1): 45–51.

23. Mez J, Mukherjee S, Thornton T, Fardo DW, Trittschuh E, Sutti S et al. The executive prominent/memory prominent spectrum in Alzheimer’s disease is highly heritable. Neurobiol Aging 2016; 41: 115–121.

24. Mukherjee S, Trittschuh E, Gibbons LE, Mackin RS, Saykin A, Crane PK et al. Dysexecutive and amnesic AD subtypes defined by single indicator and modern psychometric approaches: relationships with SNPs in ADNI. Brain Imaging Behav 2012; 6(4): 649–660.

25. Mez J, Cosentino S, Brickman AM, Huey ED, Manly JJ, Mayeux R. Dysexecutive versus amnestic Alzheimer disease subgroups: analysis of demographic, genetic, and vascular factors. Alzheimer Dis Assoc Disord 2013; 27(3): 218–225.

26. Mez J, Cosentino S, Brickman AM, Huey ED, Manly JJ, Mayeux R. Faster cognitive and functional decline in Dysexecutive versus amnestic Alzheimer’s subgroups: a longitudinal analysis of the National Alzheimer’s Coordinating Center (NACC) database. PLoS One 2013; 8(6): e65246.

27. Dubois B, Feldman HH, Jacova C, Hampel H, Molinuevo JL, Blennow K et al. Advancing research diagnostic criteria for Alzheimer’s disease: the IWG-2 criteria. Lancet Neurol 2014; 13(6): 614–629.

